# Biogenesis of NDUFS3-less complex I indicates TMEM126A/OPA7 as an assembly factor of the ND4-module

**DOI:** 10.1101/2020.10.22.350587

**Authors:** Luigi D’Angelo, Elisa Astro, Monica De Luise, Ivana Kurelac, Nikkitha Umesh-Ganesh, Shujing Ding, Ian M. Fearnley, Massimo Zeviani, Giuseppe Gasparre, Anna Maria Porcelli, Erika Fernandez-Vizarra, Luisa Iommarini

**Author notes:** Co-first authors (Luigi D’Angelo, Elisa Astro). Co-last authors (Anna Maria Porcelli, Erika Fernandez-Vizarra, Luisa Iommarini). To whom correspondence should be addressed: Luisa Iommarini, Department of Pharmacy and Biotechnology (FABIT), University of Bologna, Via Francesco Selmi 3, 40126, Bologna, Italy; Erika Fernandez-Vizarra Institute of Molecular, Cell and Systems Biology, University of Glasgow, University Avenue, Glasgow G12 8QQ, UK.

## Abstract

Complex I (CI) is the largest enzyme of the mitochondrial respiratory chain and its defects are the main cause of mitochondrial disease. To understand the mechanisms regulating the extremely intricate biogenesis of this fundamental bioenergetic machine, we analyzed the structural and functional consequences of the ablation of NDUFS3, a non-catalytic core subunit. We prove that in diverse mammalian cell types a small amount of functional CI can still be detected in the complete absence of NDUFS3. In addition, we have determined the dynamics of CI disassembly when the amount of NDUFS3 is gradually decreased. The process of degradation of the complex occurs in a hierarchical and modular fashion where the ND4-module remains stable and bound to TMEM126A. We have thus, uncovered the function of TMEM126A, the product of a disease gene causing recessive optic atrophy, as a factor necessary for the correct assembly and function of CI.

## Introduction

Human complex I (CI), NADH: ubiquinone oxidoreductase, is the largest multi-heteromeric enzyme of the mitochondrial respiratory chain (MRC), being composed of forty-five subunits encoded by both the nuclear (nDNA) and the mitochondrial DNA (mtDNA) (Hirst, 2013; Sazanov, 2015). This enzyme couples the transfer of two electrons from NADH to ubiquinone (coenzyme Q, CoQ) with the pumping of four protons from the matrix to the intermembrane space, thus contributing to the generation of the proton-motive force exploited for ATP synthesis (Hirst, 2013; Sazanov, 2015). The essential catalytic core of the enzyme is constituted by the fourteen so-called ‘core subunits’ evolutionarily conserved from bacteria to the mitochondria of higher eukaryotes (Baradaran *et al*, 2013; Zhu *et al*, 2016; Agip *et al*, 2018). The remaining thirty supernumerary subunits lack direct catalytic function but are essential for the stability and the biogenesis of the enzyme (Stroud *et al*, 2016). Supernumerary subunits are all nDNA-encoded, while seven core subunits (MT-ND1-6 and MT-ND4L) are encoded by mtDNA (Vinothkumar *et al*, 2014; Zhu *et al*, 2016). This gigantic complex has a distinctive L-shaped structure with a hydrophilic arm protruding into the matrix and a hydrophobic arm embedded in the inner mitochondrial membrane. It can be divided into three main functional modules: NADH dehydrogenase module (N-module), ubiquinone-binding module (Q-module), and proton translocating module (P-module), that is located entirely in the membrane arm (Sazanov, 2015). The transmembrane P-module contains all the mtDNA-encoded subunits and can be further divided into two proximal modules (ND1- and ND2-modules) plus two distal modules (ND4- and ND5-modules) (Guerrero-Castillo *et al*, 2017; Sánchez-Caballero *et al*, 2016; Stroud *et al*, 2016). Such modular organization reflects the enzyme evolution (Friedrich & Scheide, 2000; Moparthi & Hägerhäll, 2011), structure (Agip *et al*, 2018) and function (Hirst, 2013), as well as the pathways involved in its assembly (Guerrero-Castillo *et al*, 2017). CI biogenesis is an intricate process that takes place in a modular fashion, where each of the functional submodules are assembled and stabilized by a number of specific ‘assembly factors’, which can be chaperones or proteins involved in prosthetic group incorporation or post-translational modification of the subunits (Formosa *et al*, 2018). Each of the modules are joined together in different stages forming assembly intermediates, all eventually coming together at the end of the process to form the mature holoenzyme when assembly factors are released (Sánchez-Caballero *et al*, 2016; Guerrero-Castillo *et al*, 2017). To further complicate this scenario, the near totality of CI is found associated with complexes III (CIII) and IV (CIV), forming the respiratory supercomplexes (SCs) or ‘respirasomes’ (Acin-Perez & Enriquez, 2014; Milenkovic *et al*, 2017; Lobo-Jarne & Ugalde, 2018). The biogenesis of CI has been proposed as either being completed before interacting with CIII and CIV (Acín-Pérez *et al*, 2008; Guerrero-Castillo *et al*, 2017), or occurring in the context of the SCs (Moreno-Lastres *et al*, 2012a; Protasoni *et al*, 2020).

During CI assembly, the core subunit NDUFS3 is part of a subcomplex of about 89kDa containing NDUFS2 and NDUFA5 that appears in the early stages as the primary core for the generation of the Q-module (Vogel *et al*, 2007; Dieteren *et al*, 2012; Guerrero-Castillo *et al*, 2017). Hence, although it lacks a direct catalytic role, NDUFS3 is highly conserved and seminal for CI assembly, and missense mutations in *NDUFS3* gene cause severe encephalomyopathy, including Leigh Syndrome (LS) (Bénit *et al*, 2004; Jaokar *et al*, 2013, 3; Lou *et al*, 2018, 3; Pagniez-Mammeri *et al*, 2009). In addition, a recently developed conditional knock-out (KO) mouse model showed that the ablation of *Ndufs3* in skeletal muscle produces severe progressive myopathy and early mortality (Pereira *et al*, 2020).

Despite the great advances in the understanding of mammalian MRC biogenesis in general, and in the assembly of CI in particular, there are still many open questions concerning the hierarchy of the different components of these pathways. These questions are relevant not only to understand basic mitochondrial biology, but also the pathological mechanisms underlying CI deficiency in human disease.

We previously generated and performed a basic biochemical characterization of two different human cell lines in which the *NDUFS3* gene was knocked-out to study the impact of CI deficiency on tumor progression (Kurelac *et al*, 2019). Here, we further elucidate the role of the non-catalytic but still essential NDUFS3 subunit in CI assembly, stability and function. Our results show that progressive decrease or complete and permanent ablation of this subunit severely affects the stability of CI, but even when NFUFS3 is completely missing, yet a small fraction of CI is still able to assemble and display its NADH:ubiquinone oxidoreductase activity. In addition, we have defined the dynamics of CI disassembly upon the gradual repression of NDUFS3, showing that the distal P-modules containing MT-ND4 and MT-ND5 remain stable, while all the others are disassembled. Finally, we have identified TMEM126A/Opa7, whose function was thus far unknown, as a CI assembly factor.

## Results

### A residual fully assembled and functional respiratory CI is present upon *NDUFS3* ablation

We have reported previously that the genetic ablation of *NDUFS3* in two human cell lines, osteosarcoma 143B^−/−^ and colorectal carcinoma HCT116^−/−^, induced a severe decrease of both CI NADH dehydrogenase activity and CI-driven ATP production (Kurelac et al., 2019). However, CI (rotenone-sensitive) activity was not completely abolished despite the complete absence of NDUFS3 in 143B^−/−^ and HCT116^−/−^ (Supplementary Fig. 1A-B, Fig.1A; 13.62% and 20.31% of their relative isogenic controls, respectively). Such residual CI activity was evident in the two NDUFS3-null cell models also by In-Gel Activity (CI-IGA) in both isolated, fully assembled CI and in supercomplexes (SCs) (Fig. 1B-C and Supplementary Fig. 1C-D). This residual activity found in NDUFS3-null cells was undetectable in nDNA isogenic 143B cybrids carrying the homoplasmic frameshift mutation m.3571insC/*MT-ND1* (OS) and in mtDNA-depleted 143B Rho^0^ cells (Iommarini et al., 2018) (Fig. 1B). Moreover, it was completely abolished by blocking mitochondrial translation with 50 μg/mL chloramphenicol (CAP) (Supplementary Fig. 1E), indicating that the presence of a residual fully assembled and active CI persists despite the complete lack of NDUFS3. Moreover, CI-IGA was detectable in OS-93 cells in which 7% of wild type *MT-ND1* was able to partially rescue the disassembly of CI, indicating a high sensitivity of the method (Fig. 1B; (Iommarini et al, 2018)). *NDUFS3* ablation was accompanied by marked reduction of several CI subunits belonging to all the structural modules of the enzyme, whereas subunits belonging to CIII and CIV were not affected (Fig. 1D and Supplementary Fig. 1F). Such reduction of CI subunits was associated with the accumulation of subassembly species with positive signals for NDUFB6, NDUFB8 and NDUFA9. Nonetheless, 2D-PAGE confirmed the presence of a residual fully assembled CI in the SCs of 143B and HCT116 cells carrying the genetic ablation of NDUFS3 (Fig. 1E and Supplementary Fig. 1G). On the other hand, SC CIII_2_+IV and fully assembled CIII and CIV were neither structurally affected nor showed defects in their redox activity (Fig. 1E, Supplementary Fig. 1G and Supplementary Fig. 2A-B) and CS activity was normal as well (Supplementary Fig. 2A). We have previously shown that a residual rotenone-sensitive CI-driven ATP synthesis was found in both 143B and HCT116 cells completely lacking NDUFS3, compared to controls, while CII- and CIII-mediated ATP synthesis were not significantly affected (Kurelac *et al*, 2019). We here show that such residual CI-driven ATP production is completely abolished in NDUFS3-null 143B and HCT116 cells by the treatment with 50 μg/mL CAP (Fig. 1F). Furthermore, the presence of a residual fully assembled and functional CI was also evident in the mouse B16-F10 melanoma cell model lacking Ndufs3 (B16^−/−^), ruling out the possibility of a human-specific compensatory mechanism (Supplementary Fig. 3A-C).

**Figure 1.**
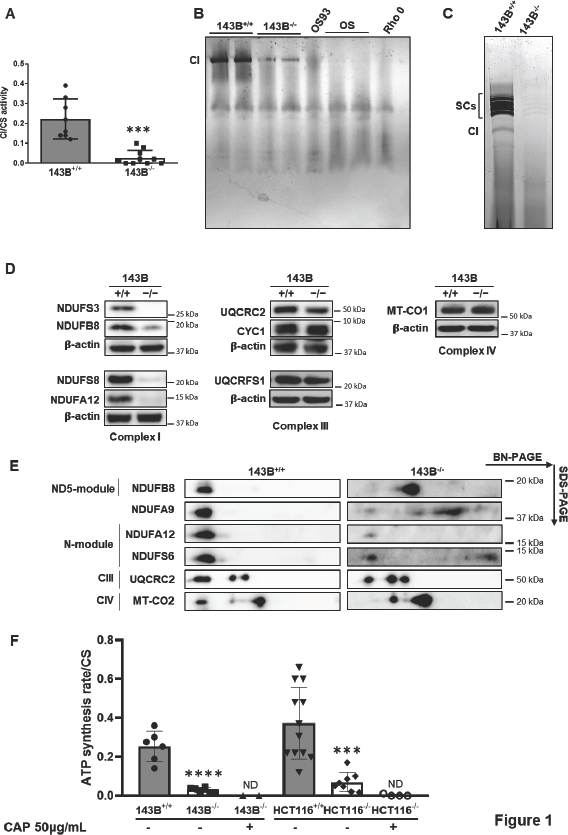
A residual, but still detectable, fully assembled and functional respiratory CI is present in constitutive *NDUFS3* KO cells. **A**) Spectrophotometric kinetic measurements of complex I (CI) activity normalized to citrate synthase (CS). Data in the graph are represented as mean ± SD (n = 9 biological replicates for 143B^−/−^ and n = 10 for 143B^+/+^). ***p = 0.0009 according to unpaired Student’s t-test. **B**) Complex I in-gel activity (CI-IGA) assay using MTT as the substrate, of 143B^+/+^, 143B^−/−^, OS93, OS and Rho0 samples solubilized with 1.6 mg DDM/mg protein and separated by High-Resolution Clear Native PAGE (hrCNE) **C**) CI-IGA assay of 143B^+/+^ and 143B^−/−^ samples, using NTB as the substrate and solubilized with 4 mg digitonin/mg protein and separated by BN-PAGE. SCs include CI+III_2_ and respirasomes (I+III_2_+IV_1-n_). **D**) Immunodetection of CI, CIII and CIV subunits on Western blots of whole-cell lysates from 143B^+/+^ and 143B^−/−^ cells lines resolved by SDS-PAGE. β-actin was used as the loading control for each blot. **E**) Immunodetection of CI, CIII and CIV subunits on Western blots of mitochondrial enriched fraction from 143B^+/+^ and 143B^−/−^ cell lines solubilized with 4 mg digitonin/mg protein and resolved by 2D BN-PAGE. All the subunits were immunodetected on the same blot. **F**) CI-driven ATP synthesis rates for 143B^+/+^ (n=6), 143B^−/−^ (n=6), 143B^−/−^ treated with 50 μg/mL Chloramphenicol (CAP) (n=3), HCT116^+/+^ (n=12), HCT116^−/−^ (n=8) and HCT116^−/−^ treated with 50 μg/mL Chloramphenicol (CAP) (n=4). Specific ATP synthesis rates (nmol/min*mg) were normalized to CS specific activity. The values represented in the graph are mean ± SD. *p < 0.0001; multiple t tests of Two-way ANOVA, Tukey’s method, with alpha=5.000%. ND: not detected.

To understand the dynamics of CI assembly upon ablation of NDUFS3, we investigated 143B cells in which the expression of NDUFS3 can be repressed by doxycycline (Dox) in a Tet-Off system, hereafter referred as 143B^−/−NDUFS3^ (Kurelac et al., 2019). Treatment with 100 ng/mL Dox induced a progressive reduction of NDUFS3 that started promptly after 12h until the subunit was virtually undetectable after 4 or 8 days of incubation (Fig. 2A and Supplementary Fig. 3D). These doses and treatment times did not affect mitochondrial translation, as shown by the normal levels of mtDNA-encoded subunit MT-CO2 (Supplementary Fig. 3E). CI-IGA was still detectable in both isolated CI and in SCs at the same time points in which the presence of NDUFS3 was undetectable (Fig. 2B and Supplementary Fig. 3F). The gradual decrease of NDUFS3 levels was not always followed by the reduction in the steady-state levels of other CI subunits, as in the case of NDUFB8, NDUFB6 and NDUFA9, which remained consistently stable after virtually complete NDUFS3 repression (Fig. 2C). Conversely, the reduction of NDUFS3 was followed by progressive decrease of fully assembled CI and accumulation of subcomplexes containing NDUFB8 and NDUFB6, while CIII and CIV were not affected (Fig. 2D-E). These data are in agreement with the presence of subcomplexes with positive immunodetection for NDUFB8 and NDUFB6 found in NDUFS3-null 143B and HCT116 cells and may explain the high levels of these subunits after NDUFS3 repression. However, a residual CI was still found in SCs after 8 days of Dox treatment of 143B^−/−NDUFS3^, as highlighted by positive staining for all the tested CI, CIII and CIV subunits (Fig. 2D-E), despite the virtually complete absence of NDUFS3. Moreover, fully assembled CI containing NDUFB8 was evident also in its isolated form after 8 days of incubation of 143B^−/−NDUFS3^ with 100 ng/mL Dox (Supplementary Fig. 3G). On the other hand, the incubation for 8 days with 100 ng/mL Dox together with 50μg/mL CAP inhibited mitochondrial translation and thus mtDNA-encoded CI subunits, resulting in the complete absence of the enzyme (Fig. 2B and Supplementary Fig. 3E). Overall, these data demonstrate that despite the complete absence of NDUFS3, either by genetic destruction or by strong expression inhibition, a residual but clearly detectable amount of active CI associated into SCs is present in both human and mouse cell models, without affecting the composition and function of the remaining OXPHOS system, whereas mtDNA-encoded subunits are an absolute requirement. The possibility that the detected CI is just residual enzyme that has not been completely disassembled can be ruled out, as treatment at the same time points with inhibitors of mitochondrial translation produced the complete disappearance of the complex (Guerrero-Castillo *et al*, 2017).

**Figure 2.**
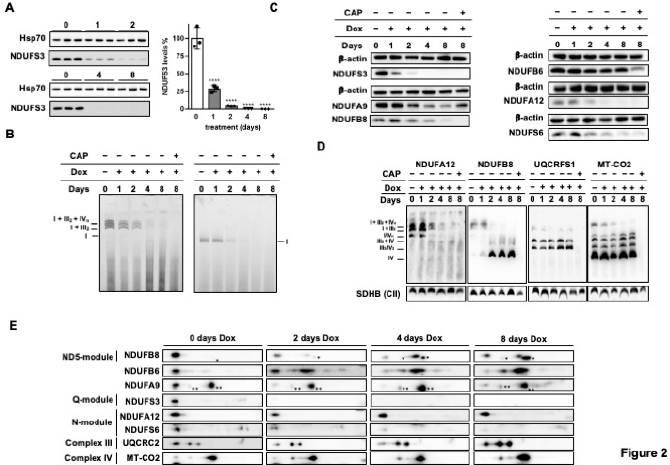
Persistence of functional respiratory CI during repression of *NDUFS3*. **A**) Immunodetection of NDUFS3 on Western blots of whole-cell lysates from 143B^−/− NDUFS3^ cells treated with 100 ng/mL doxycycline (Dox) for 0 (untreated), 1, 2, 4 and 8 days (n = 3 biological replicates). NDUFS3 band signal intensities were quantified by densitometry and normalized to the signal of Hsp70. The mean values of the treated cells were referred to those of untreated control set to 100%. Data represented on the graph are mean ± SD. ****p < 0.0001 treated vs. untreated; ANOVA with Sidak’s multiple comparisons test. **B**) Immunodetection of CI subunits on Western blots of 143B^−/−NDUFS3^ cells, treated either with 100 ng/mL for 0 (untreated), 1, 2, 4 and 8 days or simultaneously with 100 ng/mL Dox and 50 μg/mL CAP for 8 days, resolved by SDS-PAGE. β-actin was used as the loading control. **C**) CI-IGA, using NTB as the substrate, of enriched mitochondrial fraction from repressed 143B^−/−NDUFS3^, solubilized either with 4 mg digitonin/ mg protein or 1.6 mg/DDM/ mg protein and separated by BN-PAGE. SCs include CI+III_2_ and respirasomes (I+III_2_+IV_1-n_). **D**) Immunodetection of CI subunits NDUFA12 (N-module) and NDUFB8 (ND5-module), CIII subunit UQCRFS1 and CIV subunit MT-CO2, in Western blots of enriched mitochondrial fraction from 143B^−/−NDUFS3^ cells treated as in B solubilized either with 4 mg digitonin/ mg protein or 1.6 mg DDM/ mg protein and separated by BN-PAGE. SCs include CI+III_2_ and respirasomes (I+III_2_+IV_1-n_). SDHB (CII) was used as the loading control. **E**) Immunodetection of CI, CIII and CIV subunits on Western blots of mitochondrial enriched fraction from 143B^−/−NDUFS3^ cells treated with 100 ng/mL Dox for 0 (untreated), 2, 4 and 8 days, solubilized with 4 mg digitonin/mg protein and resolved by 2D BN-PAGE. All the subunits were immunodetected on the same blot. One asterisk (*) indicates a non-specific signal derived from an anti-NDUFA8 antibody with which the membrane was previously incubated. Two asterisks (**) indicate a signal derived from antibody anti-MT-CO1.

### Progressive loss of NDUFS3 reveals the modular dynamics of CI disassembly

The Tet-Off 143B^−/−NDUFS3^ model provided a dynamic system in which the kinetic of CI disassembly could be followed upon the progressive depletion of NDUFS3. Changes in mitochondrial protein levels relative to the untreated 143B^−/−NDUFS3^ cells were analyzed upon treatment with 100 ng/mL Dox after 2, 4 and 8 days by quantitative proteomics using SILAC in duplicate experiments and validated via western-blot immunodetection (Fig. 3A and Supplementary Fig. 4A). When the fold change values for each protein in the two experiments were plotted (log2 H/L ratios of one experiment in the x axis; −log2 H/L rations of the second experiment in the y axis), the protein points distributed over a 45° diagonal line, reflecting the correlation between the two duplicate experiments (Fig. 3A). Most of the detected proteins were unaffected by the progressive reduction of NDUFS3 levels, as they clustered around the axis origin. In particular, subunits of CIII and CIV were found unaltered upon treatment with 100 ng/mL Dox up to 8 days (Fig. 3A), confirming the stability of these complexes when NDUFS3 is repressed. On the other hand, significant progressive changes in the relative abundance of CI subunits and assembly factors were detected (Fig. 3A-B and Supplementary Fig. 4B). Most of the CI subunits were detected in these time course experiments, ranging from 77% to 82% of the 45 CI subunits. In agreement with western blotting data, NDUFS3 steady state levels underwent a progressive severe decrease upon treatment with Dox that reached a logarithmic fold change of −3.73±0.27 (approximately 13-fold decrease) after 8 days (Fig. 3A-B). Q-module subunits (NDUFS2, NDUFS7, NDUFS8 and NDUFA5) were the first to be downregulated after NDUFS3 depletion, being significantly reduced already after 2 days of Dox treatment (Fig. 3A-C). The levels of N-module subunits (NDUFV1, NDUFV2, NDUFV3-L, NDUFS1, NDUFS4, NDUFS6, NDUFA2, NDUFA12, NDUFA6 and NDUFA12) were less affected after 2 days of NDUFS3 suppression, but reached significant reduction after 4 and 8 days (Fig. 3A-C). Interestingly, subunits NDUFA6 and NDUFA7, whose module assignment is still debated, clustered with N-module subunits with a reduction of −2.31 ± 0.06 and −2.22 ± 0.01 respectively, suggesting an association with this module. The stability of P-module subunits was heterogeneous, as some of them were not affected by NDUFS3 repression (Fig. 3A-C). On the contrary, ND1-module subunits (ND1, NDUFA3, NDUFA8 and NDUFA13) were markedly destabilized by NDUFS3 suppression and clustered together with Q- and N-modules after 8 days of Dox treatment. Among ND1-module subunits, NDUFA3 was the most stable with a log_2_ ratio of −1.89±0.35. At the earliest time-point ND2-, ND4- and ND5-modules were virtually unaffected by NDUFS3 repression. However, subunits belonging to ND2-module (ND2, ND3, NDUFA10, NDUFS5 and NDUFC2) became more unstable after 8 days of Dox treatment. Interestingly, NDUFA10 and NDUFS5 protein levels clustered with ND1-module subunits rather than those belonging to the ND2-module, with a logarithmic fold change of −2.86±0.17 and −2.92±0.15 respectively. On the other hand, the ND4- and ND5-modules were particularly stable upon NDUFS3 depletion, with their levels unchanged at all times with respect to those of the control. The permanence of the distal membrane arm subunits was most probably due to the formation of stable CI subassemblies, as shown by the accumulation of a NDUFB6- and NDUFB8-containing subassembly intermediate (Fig. 1E, Fig. 2D-E and Supplementary Fig. 1G). NDUFA9, NDUFA11, NDUFB4 and NDUFB6 subunits are shown as a group termed “uncertain” as their module assignment is still unclear (Formosa *et al*, 2018).

**Figure 3.**
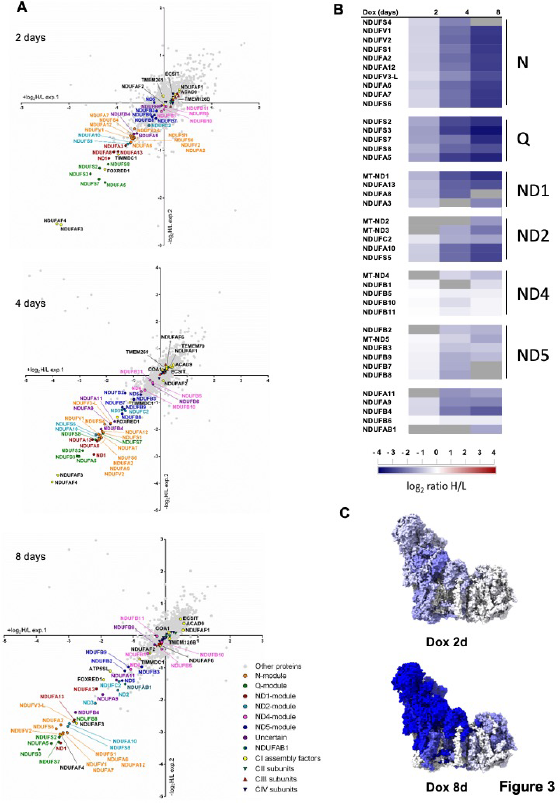
Progressive loss of NDUFS3 reveals the modular dynamics of CI disassembly. **A**) Scatter plots generated from the MS analysis of duplicate SILAC experiments (experiments 1 and 2) of comparing untreated 143B^−/−NDUFS3^ with cells treated with 100 ng/mL Dox for 2, 4 or 8 days. Mitochondrial enriched fractions for each experiment were obtained and analyzed by LC-MS. The values of the logarithmic fold change (log_2_H/L) for each protein in experiment 1 (exp.1; Light untreated, Heavy Dox treated) are represented in the x-axis. The logarithmic fold change values (-log_2_H/L) derived from experiment 2 (exp.2; Heavy untreated, Light Dox treated) are represented in the y-axis. Each point represents a specific protein. CI subunits and assembly factors are highlighted; CI subunits assigned to the same module are represented with the same color. The group termed “uncertain” includes the subunits with a still unclear domain assignment (Stroud *et al*, 2016). Proteins showing statistically significant changes are also indicated in the graphs. **B**) Heatmap generated from the mean of the duplicate SILAC experiments for the detected CI subunits at each of the Dox treatment points (2, 4 and 8 days). Grey: not detected either in one or both of the duplicate experiments. **C**) CI structural subunit relative protein abundance changes induced by repression of NDUFS3 after 2 and 8 days of Dox treatment depicted on the structure of active mouse CI (PDB: 6G2J) (Agip *et al*, 2018) using ChimeraX v1.1 (Goddard *et al*, 2018).

The abundance of most CI assembly factors was unchanged upon NDUFS3 repression (Supplementary Fig. 4A-B), with the exception of NDUFAF3, NDUFAF4, FOXRED1 and ATP5SL. Interestingly, the Q-module assembly factors NDUFAF3 and NDUFAF4 levels declined significantly already after 2-days of NDUFS3 repression and remained very low during the whole time-course. The levels of two assembly factors of the ND4-module FOXRED1 and ATP5SL (Fassone *et al*, 2010; Formosa *et al*, 2015; Guerrero-Castillo *et al*, 2017; Stroud *et al*, 2016) were also reduced to a lesser extent in NDUFS3-repressed cells, in contrast with those of the structural subunits of the same module, which were consistently unchanged. Therefore, these assembly factors may belong to another module(s) or have a role in the joining of the ND4-module with the others. These data depict a dynamic scenario which indicates that CI disassembly is also a modular process in which the Q-module is promptly affected by the reduced availability of NDUFS3, followed by the nearby N-module and proximal P-module, while levels of the subunits belonging to the distal ND4- and ND5-modules are virtually unaffected and accumulate in stable assembly intermediates most likely containing NDUFB6 and NDUFB8.

### TMEM126A is a novel assembly factor of respiratory CI

The accumulation of such assembly intermediates prompted us to search for proteins that would be upregulated upon NDUFS3 repression and likely to interact with the accumulated ND4- and ND5-modules. Such proteins are *bona fide* candidates to be CI assembly factors. To this end, CI was immunopurified from 143B^−/−NDUFS3^ cells harvested after 2, 4 and 8 days of NDUFS3 repression and quantified by SILAC (Supplementary Fig. 5A). The subunit coverage detected in these experiments ranged from 40 to 43 out of 44 different CI subunits. Among all the proteins preferentially enriched upon NDUFS3 repression, TMEM126A/Opa7 was the only one showing significantly increased association with CI after 4 and 8 days of Dox treatment (Supplementary Fig. 5A). The accumulation of TMEM126A in CI enriched fractions was not mirrored by a general increase of the protein in total lysates and mitochondrial fractions from Tet-Off model upon progressive repression of NDUFS3 (Fig. 3A and Fig. 4A). Indeed, TMEM126A specifically accumulated in the CI subassembly containing NDUFB6, corresponding to the stable ND4-module found in proteomics (Fig. 4B). The (up to now) uncharacterized mitochondrial protein TMEM126A is a paralog of the CI assembly factor TMEM126B (Elurbe & Huynen, 2016; Heide *et al*, 2012; Andrews *et al*, 2013) and mutations in *TMEM126A* (OMIM, **#**612988) have been found in multiple families affected by non-syndromic autosomal recessive optic atrophy (arOA) (OMIM, #612989). However, its function and putative role in mitochondrial respiratory chain biogenesis has never been unveiled, despite its hypothetical role as a CI assembly factor candidate and its relevance in human disease. To decipher the role of TMEM126A on respiratory chain biogenesis, we investigated CI assembly and function in commercially available HAP1 cells in which *TMEM126A* was knocked out by CRISPR/Cas9 (HAP1^−/−^, Horizon Discovery, #HZGHC05796c002). In these cells, the lack of TMEM126A was associated with a reduction of the steady state levels of several subunits of CI, while those belonging to CIII and CIV were not affected (Fig. 4C-D). From a functional point of view, rotenone-sensitive CI activity was strongly reduced in HAP1^−/−^ compared to controls (Fig. 4E and Supplementary Fig. 5B), whereas CIII and CIV activities were not affected (Supplementary Fig. 5C), indicating that loss of TMEM126A induced an isolated, specific CI deficiency. Such a functional defect of CI was also translated into a reduced respiratory capacity in the HAP1^−/−^ cells (Fig. 4F). Accordingly, HAP1^−/−^ were not able to survive under metabolic stress conditions, i.e. 5 mM galactose as carbon source, and the same cells showed reduced proliferation when grown in medium containing 25 mM glucose (Fig. 4G and Supplementary Fig. 5D-E). Overall, these data demonstrate that TMEM126A is responsible for the maintenance of a functional OXPHOS system, acting as an assembly factor necessary for the correct biogenesis of CI, likely through interaction with the ND4-module, allowing this module to join the other ones to complete the assembly.

**Figure 4.**
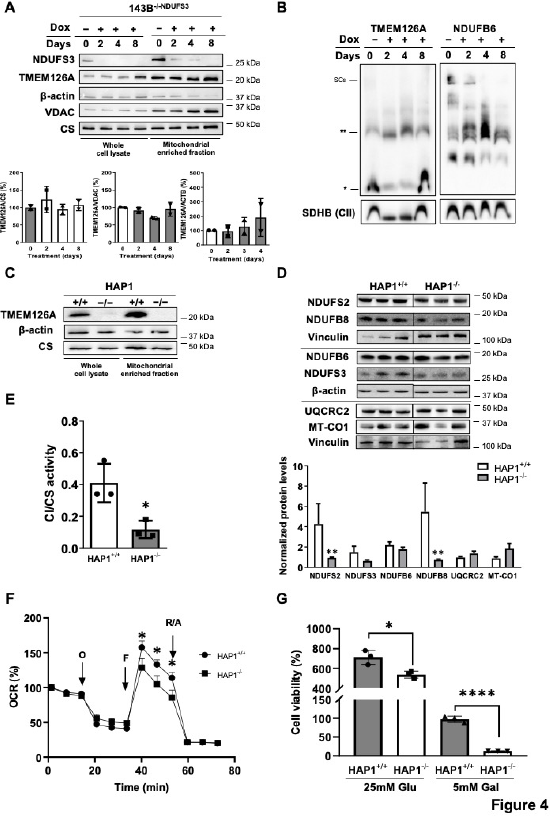
TMEM126A is required for a complete biogenesis of respiratory CI. **A**) Immunodetection of TMEM126A and NDUFS3 on Western blots of whole-cell lysates and mitochondrial enriched fractions from 143B^−/−NDUFS3^ cells treated with 100 ng/mL doxycycline (Dox) for 0 (untreated), 1, 2, 4 and 8 days (n = 2). TMEM126A band signal intensities were quantified by densitometry and normalized to the signal of either vinculin for whole cell lysates and citrate synthase (CS) or voltage-dependent anion channel (VDAC) for mitochondrial enriched fractions. The mean values of the treated cells were referred to those of untreated control set to 100%. Data were represented on the graph are mean ± SD. One-way ANOVA and one-sample t-test were used for statistics. **B**) Immunodetection of TMEM126A and NDUFB6 (ND4-module), in Western blots of enriched mitochondrial fraction from 143B^−/−NDUFS3^ cells treated as in A solubilized 4 mg digitonin/ mg protein and separated by BN-PAGE. Two asterisks (**) indicate a subassembly containing TMEM126A and co-migrating with NDUFB6 subassembly; One asterisk (*) indicates a lower molecular sized subassembly with positive staining for TMEM126A. SDHA (CII) was used as the loading control. **C**) Immunodetection of TMEM126A on Western blots of whole-cell lysates and mitochondrial enriched fractions from HAP1^+/+^ and HAP1^−/−^ cells. β-actin and CS were used as the loading controls. **D**) Immunodetection of CI, CIII and CIV subunits on Western blots of whole-cell lysates from HAP1^+/+^ and HAP1^−/−^ cells lines (n = 3-4 biological replicates) resolved by SDS-PAGE. β-actin or vinculin were used as the loading controls. Band signal intensities were quantified by densitometry and normalized to the signal of the relative loading control. The mean values of the treated cells were referred to those of untreated control set to 100%. Data were represented on the graph are mean ± SD. **p < 0.01 according to unpaired Multiple t-test, Holm-Sidak method, with alpha = 0.05. **E**) Spectrophotometric kinetic measurements of CI activity normalized to citrate synthase (CS). Data in the graph are represented as mean ± SD (n = 3 biological replicates) ***p = 0.0189 according to unpaired Student’s t-test. **F**) Oxygen consumption rate (OCR) profile in HAP1^+/+^ and HAP1^−/−^ cells. Measurements of OCR were performed upon injection of 1 μM oligomycin (O), 0.25 μM FCCP (F), 1 μM rotenone plus antimycin A (R/A) diluted in Seahorse medium, pH 7.4. Data are expressed as percentage referred to initial basal respiration and normalized on SRB absorbance and shown as mean+SEM of n = 4 independent biological replicates, *p<0.01 according to Multiple t-test Holm-Sidak method, with alpha = 0.05. **F**) HAP1^+/+^ and HAP1^−/−^ cell viability determined by sulforhodamine B (SRB) assay after 72 hours of incubation with either 25 mM glucose or glucose-free medium containing 5mM galactose. Data are mean ± SD of n = 4 independent biological replicates, *p= 0.0197 and ***p<0.0001 according to two-tailed unpaired Student’s t-test.

## Discussion

In this study, we have investigated the role of the core subunit NDUFS3 in the assembly, stability and function of respiratory CI. By exploiting multiple unique cell models where we have knocked out or repressed NDUFS3 selectively, we demonstrate that a small portion of fully assembled and functional respiratory CI is still present at the level of SCs also when NDUFS3 is absent. NDUFS3 is a core subunit, highly conserved across evolution and believed to be essential for CI assembly, being an early-entry of the Q-module (Formosa *et al*, 2018; Sánchez-Caballero *et al*, 2016; Vogel *et al*, 2007). However, although the lack of NDUFS3 strongly affects the amount of fully assembled and functional CI, a residual activity was evident in our models derived from three different human and murine cancer cell lines. It is important to note that the existence of residual CI was evident also in differentiated mouse skeletal muscle tissue of a conditional *Ndufs3* KO model (Pereira *et al*, 2020), indicating that such phenotype is not a prerogative of cancer cells. In particular, the presence of the N-module subunits at the levels of SCs indicated completion of active CI assembly. Our hypothesis is that an alternative, although not prominent, route for CI assembly can be followed when NDUFS3 is missing. SCs may help stabilize and activate the NDUFS3-less CI, providing a platform for the stabilization of mature CI as also suggested by others (Moreno-Lastres *et al*, 2012a; Protasoni *et al*, 2020). Whether this alternative route can be followed also when other CI subunits are missing or is exclusive to NDUFS3 warrants future work, which is currently underway. Indeed, NDUFS3 has the peculiarity of being a hydrophilic core subunit without real catalytic activity and when it is missing, several other subunits are reduced but not completely absent, supporting the possibility for some residual CI to complete the assembly and become functionally active. Unsurprisingly, we have shown that mtDNA-encoded subunits are pivotal for the assembly of the enzyme and no residual dehydrogenase activity was detected in the presence of a truncative mutation of *MT-ND1* or when mtDNA was depleted (Rho0 cells). In addition, no residual CI-associated dehydrogenase activity was detected when mitochondrial translation was inhibited by CAP. On the other hand, the ablation of accessory subunits was shown to impact on CI assembly to a different extent depending on the role and topology of the subunit (Stroud *et al*, 2016).

The NDUFS3 Tet-Off system provided an extremely valuable tool for the analysis of CI disassembly and for the assignment of yet uncertain subunits to specific modules. Upon NDUFS3 repression, an intact CI remains stable for about two days and then starts to decrease, indicating that a complex turnover takes place within this timeframe (Dieteren *et al*, 2012). The kinetics of the reduction of CI subunit amounts reported here is in agreement with the current CI organization and modular assembly models (Guerrero-Castillo *et al*, 2017; Stroud *et al*, 2016). Q-module subunits were significantly reduced after only 2 days of NDUFS3 repression, followed by N-module subunits, confirming that the assembly of the N-module is independent from the Q-module, but also that their interaction is necessary to build the whole CI. Regarding the membrane arm modules, the most affected subunits are those belonging to the proximal P-module (ND1 and ND2 submodules), while the distal P-module’s subunits (ND4- and ND5-modules) remain stable and form a subcomplex of about 350 kDa containing NDUFB6 and NDUFB8 which progressively accumulates with the disassembly of the fully assembled enzyme. The pattern of steady state levels of individual subunits may help the process of module assignment for some subunits whose localization is still uncertain. For example, our data support the assignment of NDUFA6 and NDUFA7 to the N-module, as proposed by Stroud et al. and in contrast with the most recent model (Guerrero-Castillo *et al*, 2017; Stroud *et al*, 2016). Likewise, abundance changes of NDUFB6 clustered with those of ND4-module subunits and remained consistently stable across the different time points, suggesting its assignment to this module. Interestingly, ND2-module subunits NDUFA10 and NDUFS5 behaved differently relative to the other module subunits being markedly affected by NDUFS3 repression. This discrepancy was observed across the time-points and their fold changes were similar to those of N-module subunits. Since NDUFA10 and NDUFS5 form extensive interactions with Q- and ND1-modules, one possibility is that the assembly impairment of such modules could be transferred to these subunits in case they are added to a preformed ND2-module.

The lack of NDUFS3 and the consequent decrease of CI amount and activity did not affect the stability or activity of other MRC complexes, in particular CIII_2_ and CIV. This is a common phenomenon for mutations in structural or ancillary CI proteins, where the biochemical manifestation is always isolated CI deficiency (Ghezzi & Zeviani, 2018). Contrariwise, mutations causing a strong impairment in CIII_2_ biogenesis are often associated with secondary loss of CI and, occasionally, CIV activities (Acín-Pérez *et al*, 2004; Barel *et al*, 2008; Carossa *et al*, 2014; Feichtinger *et al*, 2017; Lamantea *et al*, 2002; Tucker *et al*, 2013). In some cases, strong defects in CIV biogenesis also have an impact on CI assembly or stability (Diaz *et al*, 2006). In human cells, the near totality (90-95%) of CI appears associated with SCs when the mitochondria are solubilized with a mild detergent such as digitonin, whereas approximately 50% of CIII_2_ is inside SCs and 90% of CIV is in the ‘free’ monomeric form (Lobo-Jarne & Ugalde, 2018). There is growing evidence in the literature that show alternative maturation routes for CIII_2_ and CIV, depending on whether they are in the free or in the SC-associated forms (Lobo-Jarne *et al*, 2020; Moreno-Lastres *et al*, 2012b; Timón-Gómez *et al*, 2020). Moreover, only a residual amount of CI is able to mature when CIII_2_ is absent (Protasoni *et al*, 2020), therefore, the efficient incorporation of the N-module of CI only occurs in the context of the SCs, at least in human cells (Moreno-Lastres *et al*, 2012b; Protasoni *et al*, 2020). The dependent assembly of CI on one hand, and independent biogenesis of CIII_2_ and CIV on the other, would explain why defects in the latter induce secondary defects in CI assembly but not the opposite. However, this phenomenon was also viewed as the consequence of an oxidative damage-triggered degradation, by yet undescribed mechanisms, of fully assembled CI if CI is not associated with CIII_2_ (Acín-Pérez *et al*, 2004; Guarás *et al*, 2016).

Importantly, we found that the mitochondrial protein TMEM126A/Opa7 is consistently accumulated with distal P-module subcomplexes upon NDUFS3 repression. *TMEM126A* encodes an evolutionary conserved transmembrane mitochondrial protein of yet unknown function. TMEM126A shows high sequence homology with the assembly factor TMEM126B, which is part of the mitochondrial CI intermediate assembly (MCIA) complex, necessary for the assembly of the ND2-module (Formosa *et al*, 2020; Heide *et al*, 2012). The homozygous nonsense c.163C>T founder mutation (p.Arg55X) in *TMEM126A* is pathogenic in both syndromic and non-syndromic forms of autosomal recessive optic atrophy (arOA) (Désir *et al*, 2012; Hanein *et al*, 2009; La Morgia *et al*, 2019; Meyer *et al*, 2010). A partial CI deficiency and elevated lactic acid levels in blood, a typical sign of OXPHOS dysfunction, have been reported in some patients (Hanein *et al*, 2009; La Morgia *et al*, 2019). Moreover, downregulation of TMEM126A is associated with poor prognosis in breast cancer and increases metastatic properties of breast cancer cells through the activation of epithelial-to-mesenchymal transition (EMT) (Sun *et al*, 2019). Despite the considerable involvement in human pathology, TMEM126A function has so far remained unexplored. We here demonstrate that this protein is a factor necessary for the correct biogenesis and function of the mitochondrial respiratory chain, in particular of CI, through the stabilization of the submodule containing MT-ND4 within the distal P-module. Three main lines of evidence support the fact that TMEM126A is involved in this specific aspect within the CI assembly pathway. First, there is an accumulation of subunits belonging to this module inside stable subassembly species in the cells lacking NDUFS3, while other topological distinct subunits are strongly reduced. Secondly, our proteomic and interaction analysis demonstrated that TMEM126A is also accumulated in these cells and physically interacts with the remaining CI subunits. Thirdly, cells lacking TMEM126A are deficient in CI function and biogenesis.

## Conclusions

We here show that, although at low levels, fully assembled and functional CI can exist also in the complete absence of the core subunit NDUFS3. This suggests the existence of a less efficient biogenetical pathway that spares NDUFS3, thus ensuring a minimal CI activity in these conditions. This residual CI appears fully functional and able to interact with complexes III and IV to produce respiratory supercomplexes (SCs), the formation of which promotes CI assembly. In this frame, SCs may act as stabilizing factors of such residual CI. We also show that the disassembly of CI induced by progressive depletion of NDUFS3 is a modular process and revealed interactions between subunits previously undescribed. Lastly, we identify TMEM126A/Opa7 as a biogenetic factor of respiratory CI, most probably acting on the assembly of the distal P-module.

## Material and methods

### Cell lines

Human osteosarcoma 143B TK^-^ and colorectal cancer HCT116 cell lines bearing the homozygous c.9_10insCGGCG frameshift mutation in *NDUFS3*, which resulted in a premature stop codon and the truncation of the protein (p.Val6Alafs*3), were analyzed relative to their isogenic control (143B^+/+^ and HCT116^+/+^). These NDUFS3 knockout cells, referred as 143B^−/−^ and HCT116^−/−^, were generated as previously reported (Kurelac *et al*, 2019). 143B Tk^-^ cybrid cells carrying the truncative frameshift m.3571insC/*MT-ND1* mutation were also used. Specifically, the previously characterized homoplasmic cells (OS) and heteroplasmic cells with 93% of mutated *MT-ND1* (OS-93) were used (Porcelli *et al*, 2010; Gasparre *et al*, 2011; Iommarini *et al*, 2018). 143B Tk^-^ mtDNA-depleted cells (ρ0) provided CI-negative control. Doxycycline-inducible NDUFS3 knockout 143B cells (referred as 143B^−/− NDUFS3^) were also used. They were generated re-expressing *NDUFS3* using the Tet-Off inducible system ensuring the transcriptional repression of the complemented gene in the presence of doxycycline as previously described (Kurelac *et al*, 2019).

B16-F10 murine melanoma cell lines were purchased from ATCC (#CRL-6475). Cells were cultivated in Dulbecco’s modified Eagle medium (DMEM) High Glucose (Euroclone #ECM0749L), supplemented with 10% FBS (Euroclone #ECS0180L), L-glutamine (2 mM, Euroclone #ECB3000D), penicillin/streptomycin (1x, Euroclone #ECB3001D) and uridine (50 μg/mL, Sigma-Aldrich #U3003) in an incubator with a humidified atmosphere at 5% CO2 and 37°C.

Human chronic myeloid leukemia HAP1 cell line with a 79 bp-insertion in *TMEM126A* exon 4 (HAP1^−/−^; Horizon Discovery, #HZGHC05796c002) and their relative wild-type control (HAP1^+/+^) were used to demonstrate the role of TMEM126A.

### Genome editing

CRISPR/Cas9 system was used to introduce a frameshift mutation in *Ndufs3* gene in B16-F10 murine melanoma cell lines. In detail, Cas9 protein (Invitrogen #A36497) was transfected by following manufacturer’s instructions using Lipofectamine™ CRISPRMAX™ Cas9 Transfection Reagent (Invitrogen #CMAX00015) together with synthetic RNA guides designed by Deskgen and purchased from Synthego. Exon 1 targeting guide ACATGGCGGCGGCTGCAGCC with PAM sequence AGG was used to create clone E6(1) and Exon 3 targeting guide TTGTGGGTCACATCACTCCG with PAM sequence GGG was used to create G4(2). Cells were split 48 hours after transfection and DNA was extracted using Mammalian Genomic DNA Miniprep Kit (Sigma-Aldrich #G1N350). Non-homologous repair efficiency was evaluated by Sanger Sequencing using KAPA2G Taq polymerase (Kapa Biosystems #KK5601) and Big Dye protocol (Life Technologies #4337451). In particular, for Exon 1 58°C annealing temperature was used for the PCR reaction, with primers forward TGCGTCTTCTTCTCTTCGGC and reverse CAACGAAAGGCCCCAGCTAA. For Exon 3 61°C annealing temperature was used for the PCR reaction, with primers forward CTGTAACTCCAGTCTCAGGGA and reverse CACACTGCAGGGATCACTTG. Manual clonal selection was performed in order to identify the cells with frameshift *Ndufs3* mutations, leading to the generation of a pool of clones carrying the homozygous c.328A>G and c.330_331InsCT mutations. DNA extraction from 96-well plates was performed using 8 μl of Lysis Solution (Sigma-Aldrich #L3289) and 80 μl of Neutralization Buffer (Sigma-Aldrich #N9784) per sample, following manufacturer’s instructions.

### Cell maintenance and treatments

143B, HCT116 and B16-F10 cell lines were grown in high-glucose Dulbecco’s Modified Eagle’s Medium (DMEM) plus Glutamax and sodium pyruvate (Gibco-Thermo Fisher Scientific, #31966) supplemented with 10% fetal bovine serum (Gibco-Thermo Fisher Scientific, #10270), and 50 μg/mL uridine (Sigma-Aldrich, #U3003), in an incubator at 37°C with an humidified atmosphere at 5% CO_2_. 143B^−/−NDUFS3^ cells were grown in the same medium but with addition of 100 μg/mL G418 (Sigma-Aldrich, # A1720) and 50 μg/mL puromycin (Thermo Fisher Scientific, #A1113803) to sustain the selective pressure. In this cell model, expression of NDUFS3 was repressed by addition of 100 ng/mL doxycycline (Sigma-Aldrich, #D9891). For the inhibition of mitochondrial translation 50 μg/mL chloramphenicol (Fisiopharm, #J01BA01) was used. HAP1 cell lines were cultured in Iscove’s Modified Dulbecco’s Medium (IMDM) with 10% FBS, 1% Pen/Strep, and 50 μg/mL uridine. For SILAC experiments, 143B^−/−NDUFS3^ cells were grown in SILAC DMEM (Gibco Thermo Fisher Scientific, #88364) added with either isotopically labelled 152.1 mg/L L-Lysine-^15^N_2_^13^C_6_ (Sigma-Aldrich, #608041) plus 87.8 mg/L L-Arginine-^15^N_4_^13^C_6_ (Sigma-Aldrich, #608041) (“Heavy” medium), or unlabelled 143.8 mg/L L-Lysine (Sigma-Aldrich, #L8662) and 69.3 mg/L LArginine (Sigma-Aldrich, #A8094) (“Light” medium). Both differentially labelled media were supplemented also with 10% dialyzed serum (Invitrogen, #26400-044), 50 μg/mL uridine, 100 μg/mL G418, 50 μg/mL puromycin and 200 mg/mL L-Proline (Sigma-Aldrich, #P5607). Incorporation of the “Heavy” amino acids into the cells proteins was completed after approximately 9 passages (Ong *et al*, 2002). Cell lines were regularly tested for mycoplasma.

### SDS-PAGE

Whole-cell lysates were obtained incubating cell pellets with a Lysis Buffer (2% *n*-dodecyl β-dmaltoside (DDM) (Thermo Scientific, #89903) plus 1X protease inhibitor cocktail (Roche, # 11697498001) in PBS) for 20 min at 4°C. Protein concentration in the extracts was measured with a Nanodrop UV-visible spectrophotometer at λ= 280nm. 25 μg of total protein were combined with 2X Laemmli Sample Buffer (2% sodium dodecyl sulphate (SDS), 4.6% β-mercaptoethanol, 20% glycerol, 0.025% bromophenol blue and 62.5 mM Tris/HCl at pH 6.8) in one-to-one ratio, loaded on pre-cast NuPAGE^TM^ 4-12% or 12% Bis-Tris gels (Invitrogen, #NP0321, #NP0322, #NP0341, #NP0342) and resolved in a electrophoresis cell at 100 V for approximately 2 hours at RT. Either NuPAGE^TM^ MOPS or MES SDS Running Buffers were used (Invitrogen, #NP0001, #NP0002). Precision Plus Protein^TM^ Standard (Bio-Rad, #1610374) was used for molecular weight (MW) estimation.

### Blue-Native PAGE

For the preparation of samples for Blue-Native PAGE, firstly mitochondrial-enriched fractions were obtained by digitonin treatment (Nijtmans *et al*, 2002). Briefly, cells were harvested by trypsinization, 5-8×10^6^ cell pellets were washed twice and resuspended in cold DPBS and incubated on ice for 10 min with 2 mg/mL digitonin (Calbiochem, #3000410-5GM). Next, cold DPBS was added and a centrifugation at 10,000 g for 10 min and 4°C was performed. The mitochondrial-enriched fractions obtained were solubilized in a Solubilization Buffer (1.5 M aminocaproic acid, 50 mM Bis-Tris/HCl at pH 7). Proteins were quantified by a detergent-compatible Lowry-based method (Bio-Rad DC Protein Assay), solubilized either with 1.6 μg DDM/μg total proteins or 4 μg digitonin/μg total proteins and incubated at 4°C for 5 min (Wittig *et al*, 2006).Insoluble material was removed by a centrifugation (18,000 g for 30 min at 4°C) and Sample Buffer (750 mM aminocaproic acid, 50 mM Bis-Tris/HCl pH 7, 0.5 mM EDTA and 5% Coomassie Blue G250) was added. For 1D-BN PAGE, 50-75 μg of proteins were loaded on Pre-cast Native PAGE 3-12% Bis-Tris gels (Invitrogen, #BN2011BX10) at 150 V and 4°C for 4 hours. Cathode Buffer A containing 1x NativePAGE™ Running Buffer (Invitrogen, #BN2001) plus 1x NativePAGE™ Cathode Buffer Additive (Invitrogen, #BN2002); Cathode Buffer B containing 1x NativePAGE™ Running Buffer and 0.1x NativePAGE™ Cathode Buffer Additive; and Anode Buffer consisting of 1x NativePAGE™ Running Buffer were used.

### 2D BN-PAGE

1D BN-PAGE lanes were excised from the gel and incubated with a Denaturing Solution (1% 2-mercaptoethanol, 1% SDS) at RT for 1 hour. Then, the lanes were inserted in denaturing Pre-cast NuPAGE^TM^ 4-12% Bis-Tris Gels (Invitrogen, #NP0326) in a 90° orientation and separated in either MES or MOPS SDS Running Buffer at 100 V for approximately 2 hours at RT.

### Western blotting and immunodetection

Proteins resolved by either SDS-PAGE or 2D BN-PAGE were transferred onto a polyvinylidene difluoride (PVDF) membrane. The blotting was performed at 100 V for 1 hour at 4 °C in Tris-Glycine Transfer Buffer (12.5 mM Tris, 96 mM glycine, 20% Methanol, 0.025% SDS). Membranes were blocked 1 hour at RT with 5% powdered skimmed milk in PBS containing 0,1 % Tween 20 (PBS-T), washed three times for 10 min in PBS-T and incubated overnight at 4°C with the following primary antibodies diluted in a blocking solution containing 3% BSA in PBS-T: anti-TMEM126A (Atlas,#HPA046648); anti-CS (Abcam, #ab96600) 1:1000; anti-VDAC (Abcam, #ab154856) 1:1000; anti-MTCO1 (Abcam, #ab14705) 1:1000; anti-MT-CO2 (Abcam, #ab110258) 1:1000; anti-UQCRC2 (Abcam, #ab14745) 1:1000; anti-CYC1 (Proteintech, #10242-1-AP) 1:1000; anti-UQCRFS1 (Abcam, #ab14746) 1:1000; anti-NDUFA9 (Abcam, #ab14713) 1:1000; anti-NDUFS3 (Abcam, #ab110246) 1:1000 or 1:500; anti-NDUFS8 (Santa Cruz Biotechnology, #SC-515527) 1:1000; anti-NDUFB8 (Abcam, #ab110242) 1:1000; anti-NDUFB11 (Proteintech, #16720-1-AP) 1:700; anti-NDUFA12 (Sigma-Aldrich, #SAB2701046) 1:1000; anti-NDUFB6 (Abcam, #ab110244) 1:1000; anti-NDUFS6 (Abcam, #ab195807) 1:1000; anti-SDHB (Abcam, #ab147114) 1:1000; anti-SDHA (Abcam, #ab14715) 1:1000; anti-Hsp70 (Abcam, #ab2787) 1:1000; anti-β-actin (Sigma-Aldrich, #AI978) 1:1000; anti-β-tubulin (Sigma-Aldrich, #T5201) 1:1000; anti-TIMMDC1 (homemade, a generous gift from Prof. John E. Walker’s group) 1:2500. Then, three washes of 10 min in PBS-T were performed and membranes were incubated 1 hour at RT with the following 1:5000 or 1:2500 horseradish peroxidase conjugated secondary antibodies in 1% milk in PBS-T: anti-mouse (Promega, #W402B), anti-rabbit (Promega, #W401B) and anti-chicken (Promega, #G1351). Proteins resolved by 1D BN-PAGE were electroblotted as above, except that the blotting was performed at 300 mA for 1 hour at 4°C in Bicarbonate Transfer Buffer (10 mM NaHCO_3_ and 3 mM Na_2_CO_3_). The immunoreactive bands were visualized with ECL Western Blotting Reagents (GE Healthcare, #RPN2106). When performed, band signal intensities were quantified by densitometry using ImageJ and 2-way ANOVA with Sidak’s post-hoc test was used for multiple comparison statistical analysis.

### Enzymatic activity assays

Crude mitochondria were used to assess respiratory chain enzymatic activities. A pellet of 15-20 x 10^6^ cells was suspended in ice-cold Sucrose-Mannitol Buffer (200 mM mannitol, 70 mM sucrose, 1 mM EGTA and 10 mM Tris-HCl at pH 7.6) and homogenized using a glass/teflon Potter homogenizer. The obtained sample was centrifuged at 600 g for 10 min at 4 °C to discard unbroken cells and nuclei. The resulting supernatant was centrifuged at 10,000 g for 20 min at 4 °C to separate crude mitochondria from the remaining sub-cellular fractions of the sample. CI (NADH:DB:DCIP oxidoreductase), CI+CIII (NADH:cytochrome *c* reductase), CII (succinate/DCPIP), CIII activity (cytochrome *c*/DBH2), and CIV (cytochrome *c* oxidase) activities were measured as previously described (Ghelli *et al*, 2013). Each specific activity was normalized to protein content and citrate synthase (DTNB:oxaloacetate) activity. CI-IGA assays were performed either on 1D BN-PAGE or High-Resolution Clear Native PAGE (hrCNE) gels. The latter was electrophoresed as described (Iommarini *et al*, 2018) and incubated with 2 mM Tris-Cl (pH 7.4), 0.15 mM NADH, and 2.5 mg/mL 3-(4,5-dimethylthiazol-2-yl)-2,5-diphenyltetrazolium bromide (MTT) o.n. at RT. 1D BN-PAGE gels were incubated with solution containing 0.5 M Tris/HCl (pH 7.4), 0.1 mg/mL NADH, 1mg/mL nitro tetrazolium blue (NTB) at RT overnight.

### ATP synthesis rate assay

The rate of mitochondrial ATP synthesis driven by CI, CII, and CIII was measured in digitonin-permeabilized cells treated or not with 50 μg/mL chloramphenicol as previously detailed (Ghelli *et al*, 2013). Briefly, cells (10 × 10^6^/mL) were suspended in a buffer containing 150 mM KCl, 25 mM Tris–HCl, 2 mM EDTA (ethylenediaminetetraacetic acid), 0.1% bovine serum albumin, 10 mM potassium phosphate, 0.1 mM MgCl_2_, pH 7.4, and incubated with 50 μg/mL digitonin until 90–100% of cells were positive to Trypan Blue staining. Aliquots of 3 × 10^5^ permeabilized cells were incubated in the same buffer in the presence of the adenylyl kinase inhibitor P_1_,P_5_-di(adenosine-5′) pentaphosphate (0.1 mM) and OXPHOS complexes substrates, chemiluminescence was determined as a function of time with Sirius L Tube luminometer (Titertek-Berthold, Pforzheim, Germany). The chemiluminescence signal was calibrated with 11 μM internal ATP standard after the addition of 1 μM oligomycin. 1 μM rotenone was added instead of oligomycin to determine the specific CI-driven ATP synthesis. The rates of the ATP synthesis were normalized to protein content and citrate synthase (CS) activity.

### Quantitative Proteomics

Proteomic analyses were performed on mitochondrial enriched fractions and quantified using SILAC as previously described (Andrews *et al*, 2013). In the present study, portions from each sample were retained before performing CI immunoprecipitation and directly analyzed by liquid chromatography-mass spectrometry (LC-MS). For CI co-immunopurification, mitochondrial-enriched fractions were incubated overnight at 4°C with Complex I Immunocapture Kit (Abcam; #ab109711) suspension. Tryptic peptide mixtures were resolved by Nanoscale Ultra High Performance Liquid Chromatography (nano-UPLC) in a C18 reverse phase column (75 μm x 100 mm) with a gradient of acetonitrile in 0.1% formic acid and a flow rate of 300 nL/min. The eluate was analyzed with Q Exactive™ Plus Hybrid Quadrupole-Orbitrap™ Mass Spectrometer (Thermo Fisher Scientific) operating in mass tandem (MS/MS) mode. Protein identification and quantification were performed with the MaxQuant computational platform (Cox & Mann, 2008; Cox *et al*, 2011). For the relative quantification, MaxQuant calculated the ratios between the peak intensities of heavy- and light-labelled peptide pairs (H/L ratio). Each proteomic analysis was performed in duplicate: in one experiment, the heavy-labelled Dox-treated cells were mixed, in a one-to-one ratio, with the light-labelled untreated ones; in the second experiment, the labelling was inverted. Fold changes for each protein were plotted in scatter plots with the log_2_H/L ratios of one experiment in the x axis and −log_2_H/L ratios of the second experiment in the y axis. Correlation between the two duplicate experiments was reflected by the distribution of the protein over a 45° diagonal line of the scatter plots. Statistical significances of the differences for enriched and depleted proteins in each reciprocal experiment were determined by applying the B-significance test with Perseus (Cox & Mann, 2011; Tyanova *et al*, 2016).

### Cell viability assay

HAP1 cell viability was evaluated by sulforhodamine B (SRB) assay. 30,000 cells/well were seeded in 24-well plates and incubated overnight. Then, IMDM was replaced with either glucose-free and galactose-containing (5 mM) DMEM or glucose-containing (25 mM) DMEM. After 24, 48 and 72 hours of incubation, medium was replaced with a fresh one, 10% trichloroacetic acid (TCA) was added, and the plates were incubated for 1 hour at 4°C to fix the cells. 5 washes in water were performed. When the plates were dry, proteins of the fixed cells were stained by an incubation with 0.4% SRB for 30 minutes at RT. SRB was removed and 5 washes with 1% acetic acid were carried out to eliminate the residual SRB. Finally, the SRB bound to the proteins was solubilized in 10 mM Tris and the absorbance at 560 nm was read in a Victor2 plate reader (Perkin-Elmer). Measurements were blank corrected.

### Oxygen consumption rate

Mitochondrial respiration was evaluated using the Seahorse XFe Cell Mito Stress Test Kit (Agilent #103015-100) following the manufacturer instructions. Cells were seeded (40 × 10^4^ cells/well) in 80 μL of IMDM medium into XFe96 cell culture plate and allowed to attach for 24 h. Cell culture media was replaced with Seahorse XF DMEM Medium, pH 7.4 (Agilent #103575-100). OCR was measured over a 3 min period, followed by 3 min of mixing and re-oxygenation of the medium. For Mito Stress Test, complete growth medium was replaced with 180 μL of unbuffered XF medium supplemented with 10 mM glucose pH 7.4 pre-warmed at 37°C. Cells were incubated at 37°C for 30 min to allow temperature and pH equilibration. After 6 OCR baseline measurements, 1 μM oligomycin, 0.25 μM carbonyl cyanide-p-trifluoromethoxyphenylhydrazone (FCCP), and 0.5 μM rotenone plus 0.5 μM antimycin A were sequentially added to each. Three measurements of OCR were obtained following injection of each drug and drug concentrations optimized on cell lines prior to experiments. At the end of each experiment, the medium was removed and SRB assay was performed to determine the amount of total cell proteins as described above. OCR data were normalized to total protein levels (SRB protein assay) in each well. Each cell line was represented at least four wells per experiment (n = 4 replicate experiments). Data are normalized on SRB absorbance and expressed as pmoles of O_2_ per minute.

### Statistics

GraphPad Prism v.7 (GraphPad Software Inc., San Diego, CA, USA) was used to perform statistical tests and create bar plots and graphs. Unless stated otherwise, a two-tailed unpaired Student’s t-tests assuming equal variance were performed to compare averages. For each experiment, at least three biological replicates were analyzed.

## Acknowledgments

This work was supported by University of Bologna AlmaIdea Junior grant INTACT to LI, by EU H2020 ITN Marie Curie project TRANSMIT [grant number GA722605] to AMP, by Italian Ministry of Health grant DISCO TRIP [grant number GR-2013-02356666] to GG and by Core Grant from the Medical Research Council (Grant MC_UU_00015/5) to MZ. MDL was supported by Associazione Italiana Ricerca sul Cancro (AIRC) fellowship. We thank Fondazione Dal Monte (Bologna, Italy) and Centro Studi della Barbariga (Padua, Italy) for the financial support finalized to the acquisition of instruments Seahorse XFe96 and Jasco V550 spectrophotometer, respectively.

We thank Dr. Joe Carroll and Prof. John E. Walker (Mitochondrial Biology Unit, Cambridge) for the insightful discussion on the role of TMEM126A; Ana Maria Rusu and Prof. F. Musiani (University of Bologna) for the technical suggestion in the production of Complex I structure pictures.

## Author contributions

Conceptualization, M.Z., A.M.P., E.F.V. and L.I.; Methodology, M.D.L., I.K., N.U.G., S.D., I.F., E.F.V. and L.I; Investigation, L.D.A., E.A., M.D.L., S.D. and L.I.; Formal analysis, L.D.A., E.A., S.D. and L.I.; Writing – Original Draft, L.D.A., E.A., G.G., A.M.P., E.F.V. and L.I.; Review & Editing, all the authors; Funding Acquisition, M.Z., G.G., A.M.P., E.F.V. and L.I.; Supervision, E.F.V and L.I.

