## Supplemental Figures for "Biogenesis of NDUFS3-less complex I indicates TMEM126A/OPA7 as an assembly factor of the ND4-module"

This pdf contains Supplemental Figures 1 to 5.

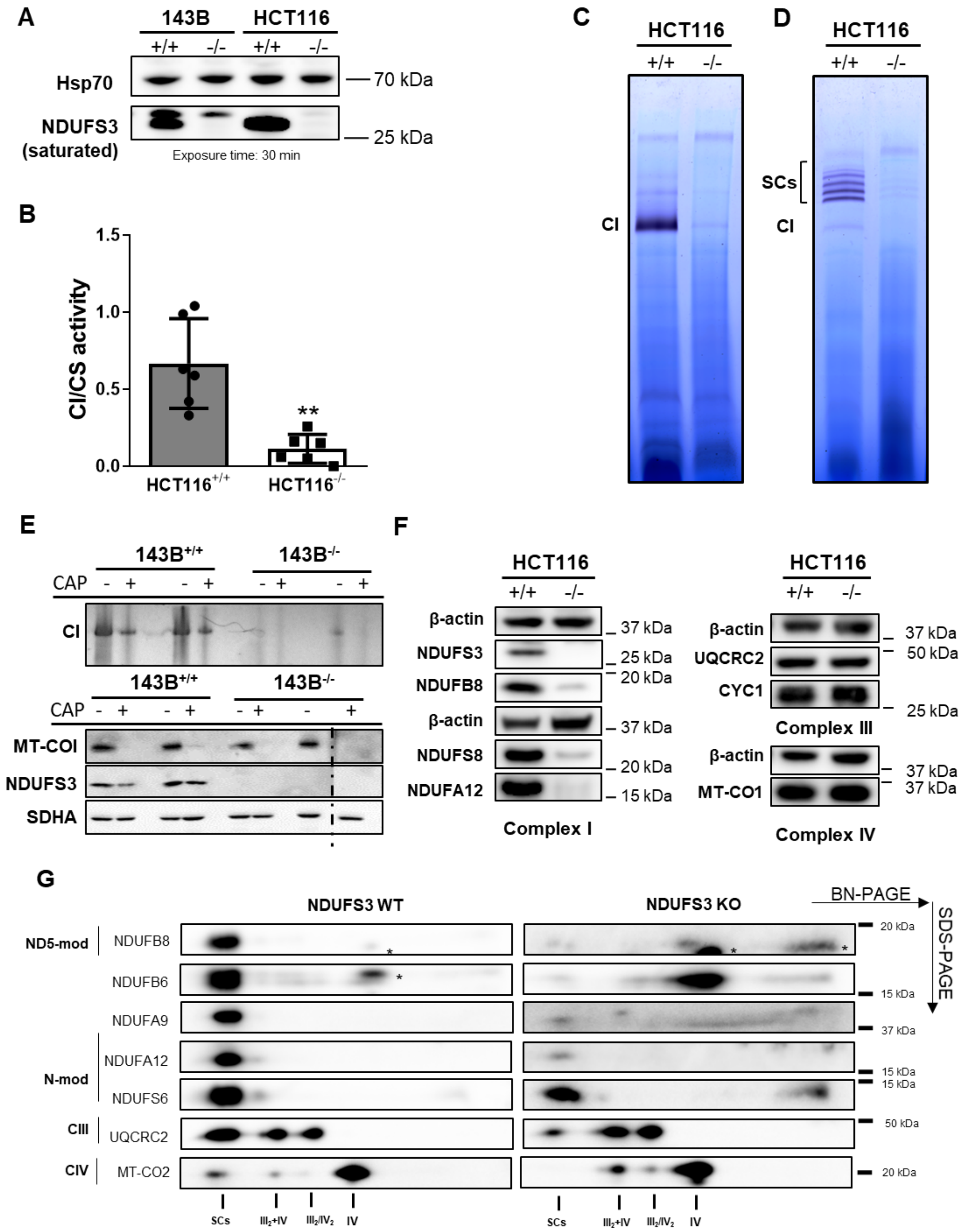


**Supplementary Figure 1**

**Supplemental Figure 1.** A residual, but still detectable, fully assembled and functional respiratory CI is present in constitutive *NDUFS3* KO cells. **A)** Immunodetection of NDUFS3 in enhancing conditions

(1:500 anti-NDUFS3 dilution, 1:2500 anti-mouse dilution, 30’ exposure) on a Western blot of whole-cell lysates from $143B^{+/+}$, $143B^{-/-}$, $HCT116^{+/+}$, $HCT116^{-/-}$ cells lines resolved by SDS-PAGE. HSP70 was used as the loading control. NDUFS3 signals in the control samples were saturated already after 2’ of exposure. **B)** Spectrophotometric kinetic measurements of CI activity normalized to CS activity for $HCT116^{+/+}$ (n = 6 biological replicates) and $143B^{-/-}$ (n = 6 biological replicates). The graphs represent mean ± SD. **p = 0.0013 according to two-tailed unpaired Student’s t-test. **C)** Complex I in-gel activity (CI-IGA) assay, using NTB as the substrate, of $HCT116^{+/+}$ and $HCT116^{-/-}$ samples solubilized with 1.6 mg DDM/mg protein and separated by BN-PAGE. **D)** Complex I in-gel activity (CI-IGA) assay, using NTB as the substrate, of $HCT116^{+/+}$ and $HCT116^{-/-}$ samples solubilized with 4 mg digitonin/ mg protein and separated by BN-PAGE. SCs include $CI+III_2$ and respirasomes ($I+III_2+IV_{1-n}$). **E)** Top: CI-IGA assay, using MTT as the substrate, of $143B^{+/+}$ and $143B^{-/-}$ samples solubilized with 1.6 mg DDM/mg protein and separated by BN-PAGE. Cells were treated (+) with 50 µg/mL chloramphenicol (CAP) for 4 days. (-) untreated. Bottom: Immunodetection of MT-CO1 and NDUFS3 on a Western blot of the same mitochondrial enriched fraction used for CI-IGA separated by SDS-PAGE. SDHA was used as the loading control. **F)** Immunodetection of CI, CIII and CIV subunits on Western blots of whole-cell lysates from $HCT116^{+/+}$ and $HCT116^{-/-}$ cells lines resolved by SDS-PAGE. β-actin was used as the loading control for each blot. **G)** Immunodetection of CI, CIII and CIV subunits on Western blots of mitochondrial enriched fraction from $HCT116^{+/+}$ and $HCT116^{-/-}$ cell lines solubilized with 4 mg digitonin/mg protein and resolved by 2D BN-PAGE. One asterisk (*) represent a non-specific signal derived from an anti-NDUFA8 antibody with which the membrane was previously incubated.

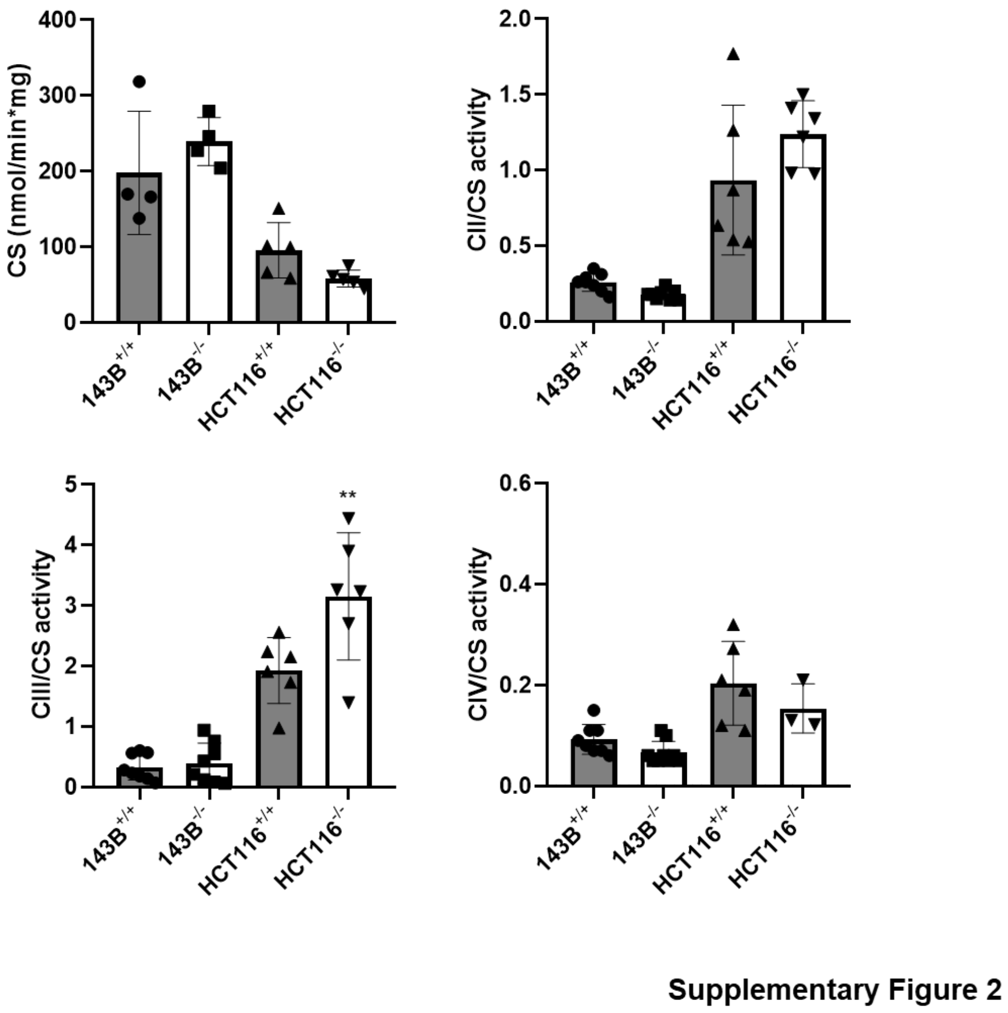


**Supplemental Figure 2.** ***NDUFS3*** **ablation does not affect the function of the rest of the respiratory chain.** Spectrophotometric kinetic measurements of citrate synthase (CS) specific activity (nmol/min*mg) and of CII, CIII and CIV activities normalized to CS activity in $143B^{+/+}$, $143B^{-/-}$, $HC116^{+/+}$, $HC116^{-/-}$ cells. Data in the graphs are represented as mean ± SD (n = 3-10 biological replicates). No statistically significant differences were found between wild type and control samples by unpaired Student's t-test, except for CIII activity in HCT116 cells (**p = 0.0065).

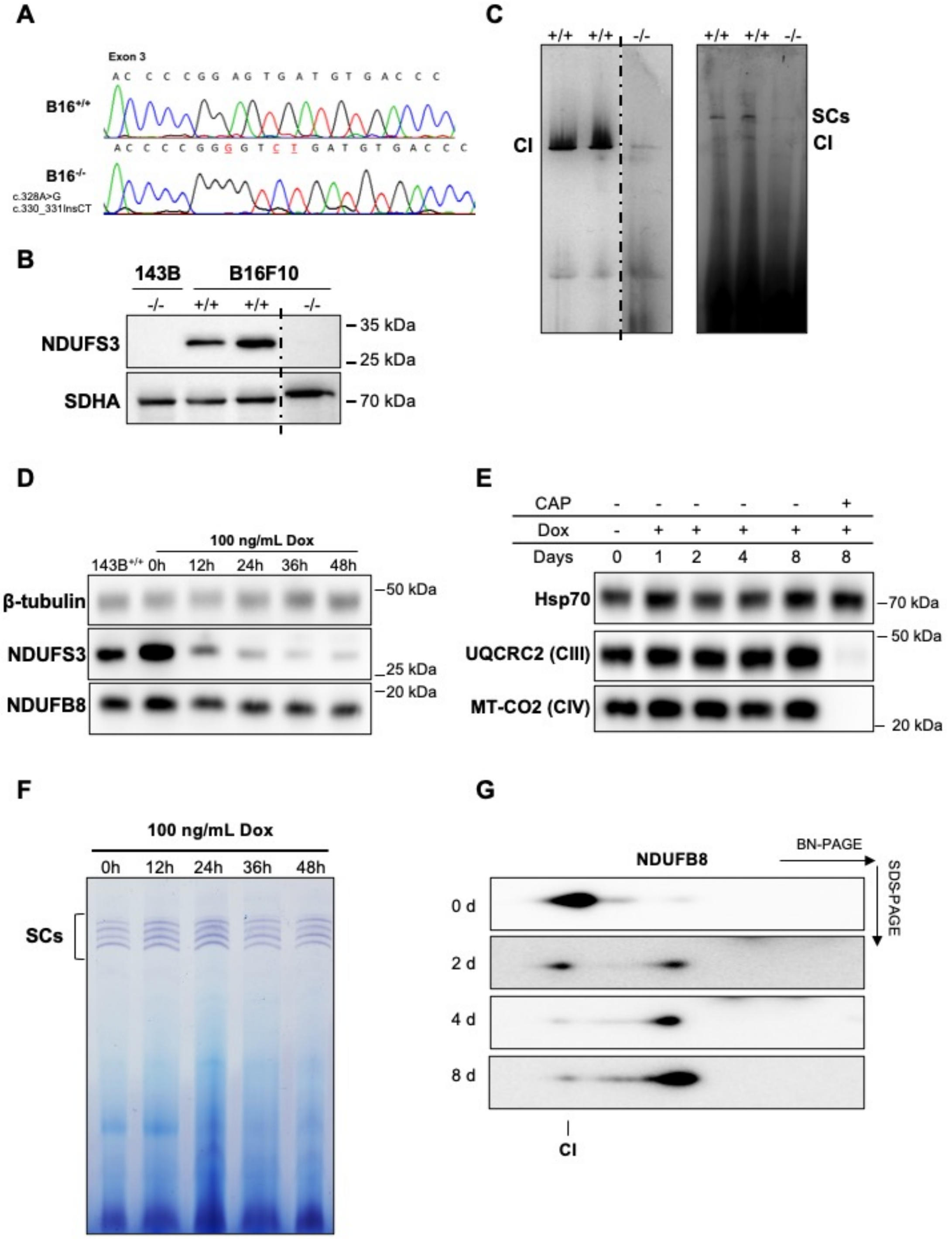


**Supplementary Figure 3**

**Supplemental Figure 3. A residual fully assembled and functional respiratory CI is present upon *NDUFS3* ablation in human and mouse derived cells. A)** Sanger sequencing-derived electropherograms showing the c.328A>G and c.330_331InsTC mutations causing Ndufs3 functional knock-out in mouse melanoma $B16^{-/-}$. **B)** Immunodetection of NDUFS3 on a Western blot of mitochondrial enriched fraction from $143B^{-/-}$ (negative control), $B16^{+/+}$ and $B16^{-/-}$ separated by SDS-PAGE. SDHA was used as the loading control. **C)** Complex I in-gel activity (CI-IGA) assay, using NTB as the substrate, of $B16^{+/+}$ and $B16^{-/-}$ samples solubilized with either 2 % digitonin (right) or 1% DDM (left) and separated by BN-PAGE. **D)** Immunodetection of NDUFS3 and NDUB8 on a Western blot of whole-cell lysates from $143B^{-/-NDUFS3}$ cells treated with 100 ng/mL Dox for 12, 24, 36 and 48 hours. Untreated $143B^{-/-NDUFS3}$ (0h) and $143B^{+/+}$ cells are the controls. β-tubulin was used as the loading control. **E)** Immunodetection of UQCRC2 and MT-CO2 on Western blots of $143B^{-/-NDUFS3}$ cells, treated either with 100 ng/mL for 1, 2, 4 and 8 days or simultaneously with 100 ng/mL Dox and 50 µg/mL CAP for 8 days, resolved by SDS-PAGE. Untreated $143B^{-/-NDUFS3}$ cells (0d) is the control. Hsp70 was used as the loading control. **F)** CI-IGA assay, using NTB as the substrate, of $143B^{-/-NDUFS3}$ treated with 100 ng/mL Dox for 12, 24, 36 and 48 hours. Samples were solubilized with 2 % digitonin and separated by BN-PAGE. **G)** Immunodetection of NDUFB8 on a Western blot of mitochondrial enriched fraction from $143B^{-/-NDUFS3}$ cells treated with 100 ng/mL Dox for 0 (untreated), 2, 4 and 8 days, solubilized with 1.6 mg DDM/mg protein and resolved by 2D BN-PAGE.

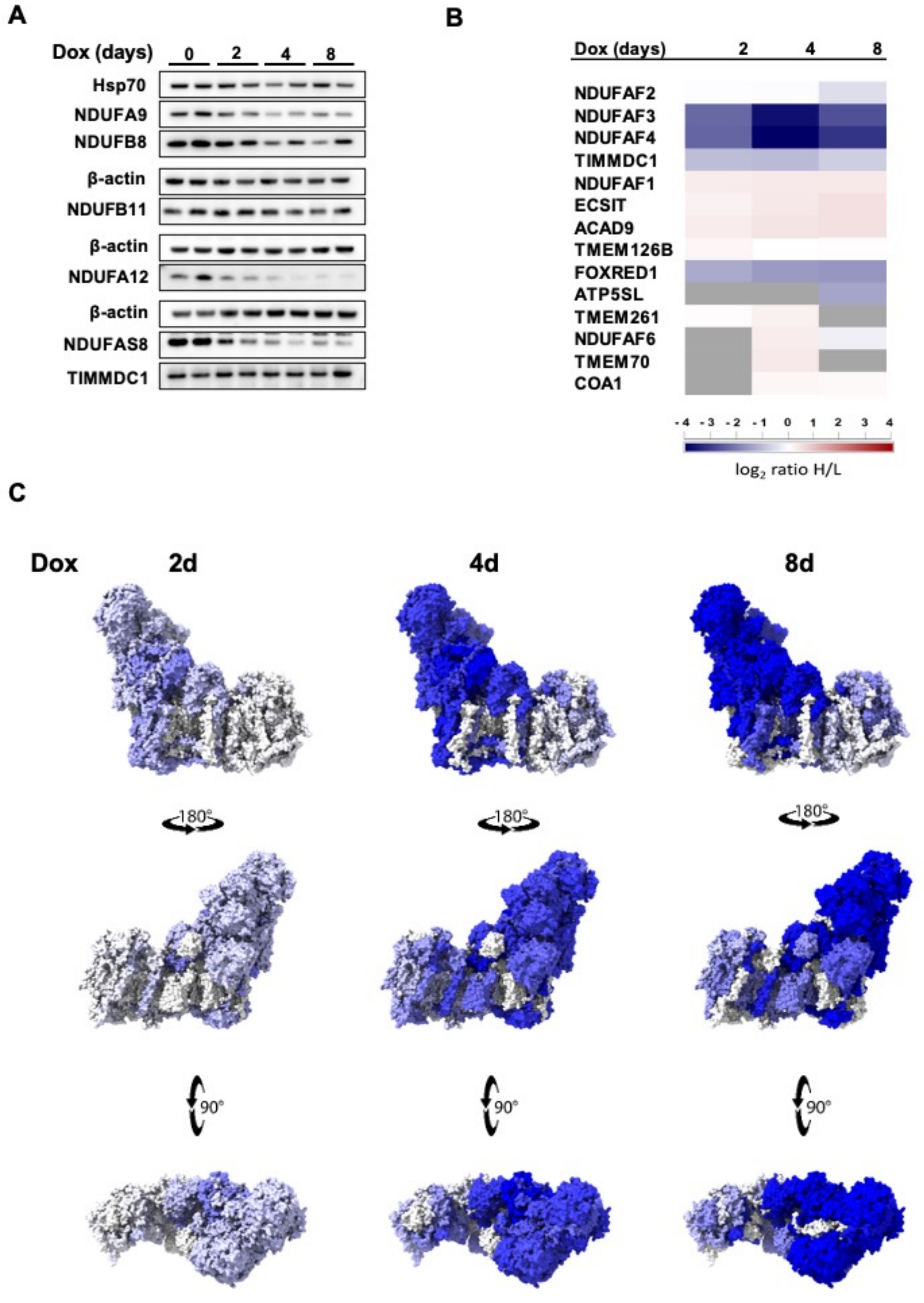


Supplementary Figure 4

**Supplemental Figure 4. Progressive loss of NDUFS3 reveals the modular dynamics of CI disassembly. A)** Immunodetection of CI subunits and CI assembly factor TIMMDC1 on Western blots of whole-cell lysates from $143B^{-/-NDUFS3}$ cells treated with 100 ng/mL Dox for 2, 4 and 8 days resolved by SDS-PAGE. Untreated $143B^{-/-NDUFS3}$ cells (0d) provided as control. Hsp70 and β-actin provided as loading controls. **B)** Heatmap generated from the mean of the two reciprocal duplicate SILAC experiments for the detected known CI assembly factors at each time point of NDUFS3 repression (2, 4 and 8 days). Grey: not detected in either one or both the duplicate experiments. **C)** CI structural subunit relative protein abundance changes induced by repression of NDUFS3 after 2 and 8 days of Dox treatment depicted on the structure of active mouse CI (PDB: 6G2J) using ChimeraX v1.1.

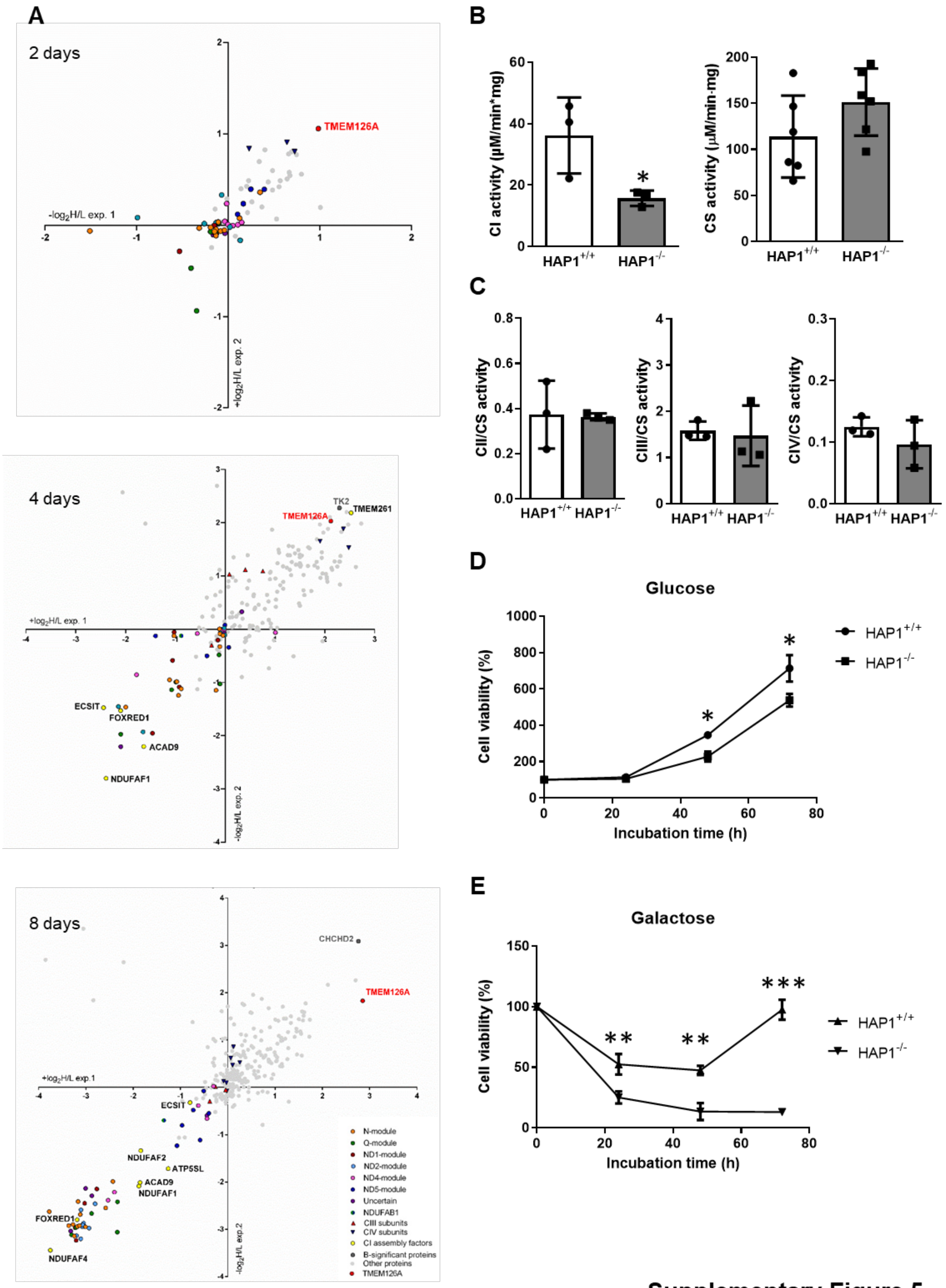


**Supplementary Figure 5**

**Supplemental Figure 5. TMEM126A is required for a complete biogenesis of respiratory CI. A)** Scatter plots generated from the MS analysis of CI immunopurification fractions performed on the same mitochondrial enriched fractions used for the proteomic analyses in Figure 3. The values of the logarithmic fold change (log2H/L) for each protein in experiment 1 (exp.1; Light untreated, Heavy Dox treated) are represented in the x-axis. The logarithmic fold change values (-log2H/L) derived from experiment 2 (exp.2; Heavy untreated, Light Dox treated) are represented in the y-axis. Each point represents a specific protein. CI subunits and assembly factors are highlighted; CI subunits assigned to the same module are represented with the same color. The group termed "uncertain" includes the subunits with a still unclear assignment (6). TMEM126A is shown in red since it is enriched in all timepoints. The proteins showing statistically significant changes are also indicated in the graphs. **B)** Spectrophotometric kinetic measurements of CI activity not normalized to citrate synthase (CS) activity (nmol/min*mg) and CS specific activity (nmol/min*mg) in $HAP1^{+/+}$ and $HAP1^{-/-}$ cells. Data in the graphs are represented as mean ± SD (n = 3 biological replicates). *p = 0.0487; unpaired Student's t-test. **C)** Spectrophotometric kinetic measurements of CII, CIII and CIV activities normalized to that of CS in $HAP1^{+/+}$ and $HAP1^{-/-}$. Data in the graphs are represented as mean ± SD (n = 3 biological replicates). No statistically significant differences were found by unpaired Student's t-test. **D-E)** Cell viability curves for $HAP1^{-/-}$ and $HAP1^{+/+}$ determined by sulforhodamine B (SRB) assay after incubation with either glucose-free and galactose-containing (5 mM) DMEM or glucose-containing (25 mM) DMEM for 24, 48 and 72 hours. Data are mean ± SD (n = 3 biological replicates), *p < 0.05, **p <0.01, ***p <0.001 according to Multiple t-test, Holm-Sidak method, with alpha = 0.05.
